# Identification and characterization of iPTH and two parathyroid hormone receptors-like (PTHR1 and PTHR2) in the tick *Ixodes ricinus*

**DOI:** 10.64898/2026.06.22.733765

**Authors:** Vanda Klöcklerová, Juraj Koči, Emma Buchová, Matej Medla, Mirko Slovák, Ladislav Roller, Dušan Žitňan

## Abstract

The tick *Ixodes ricinus* is the main vector of human and animal pathogens in Europe. Despite its importance in epidemiology and medicine, our understanding of physiological mechanisms controlling blood feeding, osmoregulation, or development are still limited. Here, we identify novel neuropeptide invertebrate parathyroid hormone-like peptide (iPTH) and its two receptors – PTHR1 and PTHR2 in *I. ricinus*. Functional aequorin-based assay confirmed specific activation of both receptors by iPTH. Using RT-qPCR we detected the PTHR1 transcript in the synganglion, while increased expression levels of PTHR2 were found in the salivary glands, hindgut and female gonads. RNA-mediated knockdown of iPTH receptors in nymphs resulted in delayed blood feeding, and a high incidence of defects in adult ecdysis. Consistent with observed phenotypes, iPTH is expressed in multiple neurons of the synganglion which project arborizing axons to the salivary glands, rectal sack and skeletal muscles. iPTH was colocalized with orcokinin-immunoreactivity (OK-IR) in all neurons that innervate these peripheral tissues. iPTH is further colocalized with tachykinin (TK) in Pd_1_DL_1_ neurons, suggesting coordinated action with other neuropeptides. Our findings indicate that iPTH signaling is required for normal feeding, development and successful ecdysis.

## Introduction

The hard tick *Ixodes ricinus* is a hematophagous ectoparasite and the most important pathogen vector in temperate Europe, which needs to be monitored by European Center for Disease Prevention and Control (ECDC, 2023). The life cycle of this tick comprises of egg and three active life stages – larva, nymph, and adult (Apanaskevich & Oliver, 2014) which needs a blood meal provided from a broad range of potential hosts to develop into the following stage (Honig et al., 2017). This process requires ticks to survive local environmental conditions in between the blood meals and process huge amounts of proteins, water and salts ingested with the blood. For this purpose, ticks evolved a very unique network of peptidergic neurons and enteroendocrine cells that control digestion and osmoregulation by the salivary glands, midgut and hindgut (Kim et al., 2016; Koči et al., 2014; Roller et al., 2015; Šimo & Park, 2014).

Parathyroid hormone (PTH) and its receptors participate in regulation of ion balance in vertebrates and their function has been intensively studied for over a century (Rendina-Ruedy & Rosen, 2022). In humans, PTH is produced by parathyroid glands and its main role is maintenance of calcium and phosphate homeostasis (Goodman & Salusky, 1996). Parathyroid hormone-like receptors (PTHRs) have been recently identified in insects and other invertebrates (Cardoso et al., 2024; Xie et al., 2020). In these animals PTHR gene underwent independent duplication, but it was lost at least twice during insect evolution (Li et al., 2013a). Moreover, invertebrate PTHRs exploit an alternative ligand with no sequence homology to conventional vertebrate PTH, named invertebrate parathyroid hormone receptor-like activaring peptide (Cardoso et al., 2024; Xie et al., 2020). Our current information on iPTH/PTHR function in arthropods are limited to a single study in the beetle *Tribolium castaneum* in which iPTH is a brain-gut neuropeptide involved in wing expansion and exoskeletal development (Xie et al., 2020). Recently, iPTH was identified in the transcriptome from the synganglion of *Rhipicephalus microplus* but there is no further information on iPTH or PTHRs in ticks (Waldman et al., 2022).

In this study, we used in silico analyses followed by molecular cloning to identify iPTH and PTHRs in the hard tick *I. ricinus*. These receptors were functionally characterized in aequorin-based *in vitro* assay in CHO cells. Utilizing visualization techniques, we described neurons producing iPTH, while molecular biology approaches were used to determine tissue- and stage-specific expression of newly identified PTHRs. Finaly, we downregulated PTHRs using RNA interference (RNAi) as a first step in elucidation of iPTH/PTHRs role in the tick physiology.

## Materials and Methods

### Experimental animals

All animal experiments were carried out in the facility of the Biomedical Research Center of the Slovak Academy of Sciences (BMC SAS) (facility permit number SK U CH 02021), in accordance with the Animal Use Protocols (permit numbers 2882/2022 and 693/2024) approved by the State Veterinary and Food Administration of the Slovak Republic.

Pathogen-free *I. ricinus* ticks originate from the breeding facility at the Institute of Zoology, Slovak Academy of Sciences. The ticks were maintained in humid chambers with 95% humidity under 10:14 h light/dark cycle at 22°C. For all experimental procedures, *I. ricinus* nymphs were fed on BALB/c mice obtained from the Institute of Experimental Pharmacology and Toxicology (CEM SAS, Dobrá voda, Slovakia), while adult female ticks were fed on white Newfoundland rabbits from the Research Institute for Animal Production (Nitra, Slovakia). Dunkin-Hartley guinea pigs used for generation of antisera to iPTH (see below) were purchased from AnLab (Czechia).

### In silico analysis and identification of putative iPTH and PTHRs sequences

Sequences of iPTH (XP_008194702.1) and PTHRs (XP_008192153.1; XP_015834182.1) of the beetle *T. castaneum* were used in BLAST search of putative *I. scapularis* iPTH and PTHRs. The resulting putative sequences (XP_029823490.2; XP_040063819.1 for iPTH, and XP_029845012.1; XP_029838018.1; XP_029844994.1; XP_029834640.1; XP_029838024.1 for PTHRs) were used in additional BLAST search in the *I. ricinus* genome database (Carpi et al., 2012; G. Carpi et al., 2011). We searched for related neuropeptide or receptor sequences using NCBI BLAST (https://blast.ncbi.nlm.nih.gov/Blast.cgi). Presence of a signal peptide in the identified sequence was confirmed using software SignalP 6.0 (https://services.healthtech.dtu.dk/service.php?SignalP). Cleavage sites in the sequence of prepropeptide were determined according to (Veenstra, 2000). Sequence alignment was made in ClustalX version 2.1 Multiple Sequence Alignment tool (http://www.clustal.org/clustal2/) and conserved domains of GPCRs were determined with NCBI CDD search tool (Lu et al., 2019).

### cDNA cloning and rapid amplification of cDNA ends (RACE)

Template cDNAs were prepared as described previously (Roller et al., 2016). Briefly, the salivary glands, gut, gonads, and CNS attached to carcass were dissected in saline buffer (140 mM NaCl, 5 mM KCl, 5 mM CaCl_2_, 4 mM NaHCO_3_, 1 mM MgCl_2_, 5 mM HEPES, pH 7.2). 5-10 individuals per observed stage and tissue were pooled into a single tube with RNA later (Sigma-Aldrich, St. Louis, MO, USA), and stored at -20°C. The following stages of ticks were used: feeding nymphs on days 1-3 (FN1-3), replete/detached nymphs on days 0, 2, 7, 14, 21, 28, 35 (RN0-35); unfed adult females (UA); feeding adults on days 1-7 (FA1-7); replete/detached adults on days 0, 1, 2, 4, 6, 8, 10 (DA0-10), ovipositing female on days 0 and 10 (OA, OA10). To distinguish iPTHR expression in different parts of the digestive system, midguts and hindguts of 10-20 unfed females were separately dissected and pooled. The synganglia attached to ventral cuticle from the unfed and feeding ticks were pooled and together with the other samples described above used for cDNA preparation. Tissues were homogenized with stainless steel beads using Tissue Lyser II (Qiagen, Cologne, Germany) and total RNA was extracted with TRI Reagent (Zymo Research, USA) or PureLink RNA Mini Kit (Invitrogen). Single-stranded cDNA was prepared with Maxima H Minus First Strand cDNA Synthesis Kit (Thermo Fisher Scientific, MA, USA) and double-stranded cDNA was prepared SMARTer cDNA synthesis kit (Takara Bio, Japan).

Verification of identified sequences was made by designing specific primers for iPTH and PTHRs. The entire ORF and parts of untranslated regions of iPTH transcript, and fragments of PTHR1 and PTHR2 ORFs were first amplified by DNA oligonucleotides (primers for iPTH – 2345+2347; PTHR1 – 2350+2417; PTHR2 – 2356+2480). Amplified iPTH, PTHR1 and PTHR2 fragments were inserted into pGEM-T Easy vector (Promega, WI, USA) and sequenced. To complete 3’ and 5’ ends of receptor sequences, we used multiple nested PCRs with SMARTer cDNA as a template, template-specific primers (1245 for 5’ends; and polyT primer 1363 for 3’ends) and receptor-specific primers. PTHR1 5’end was determined with primers 2417, 2514 in two consecutive nested PCRs in combination with primer 1245; and 3’end with three consecutive nested PCRs with primers 2530, 2377, 2396 combined with 1363. 5’ end sequence of PTHR2 was completed in two nested PCRs with primers 2384 and 2385 combined with primer 1245. Fragments of PTHRs were sequenced. The primers are listed in Table 1.

**Table 1.**
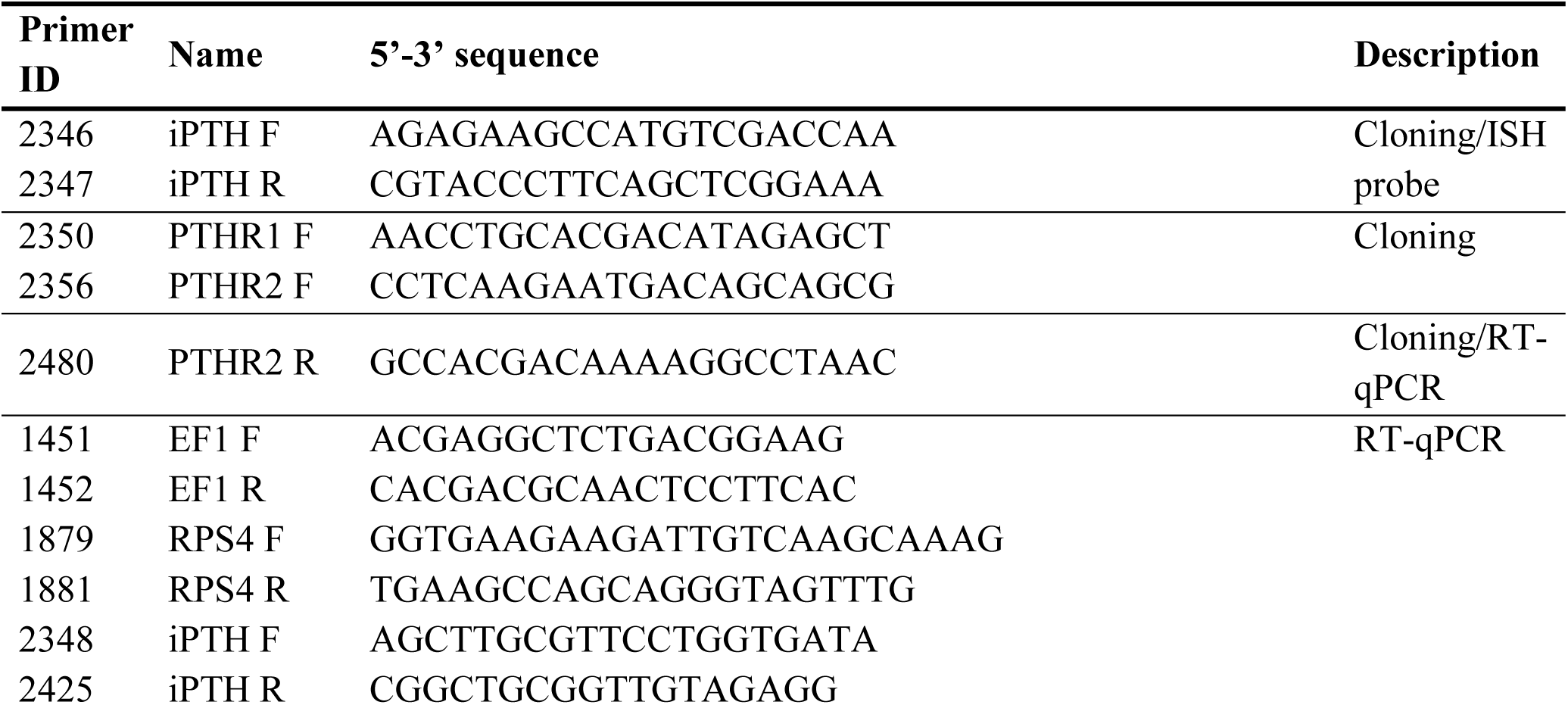
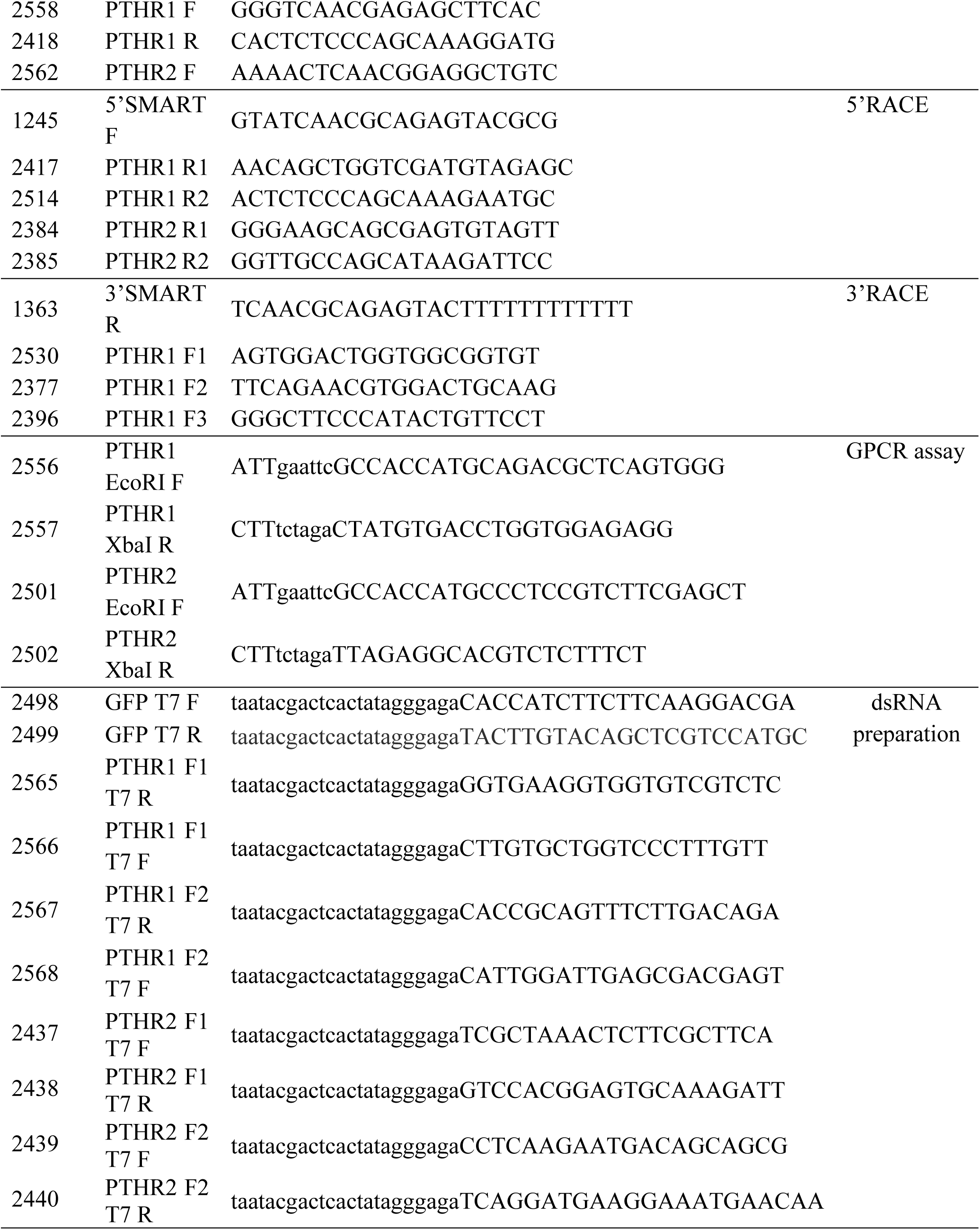
List of primers – identification number used in text, name, 5’-3’ sequence and description of each primer are listed. Recognition sites for restriction endonucleases are marked lower case in 5’-3’ sequence.

### Aequorin-based GPCR assay

Functional characterization of newly identified PTHR1 and PTHR2 receptors was done in Chinese hamster ovaries K1 cell line (CHO-K1) using aequorin-based Ca^2+^ mobilization assay as previously described (Daubnerová et al., 2021; Medla et al., 2023). Briefly, whole ORF of PTHR1 (2557; 2558) and PTHR2 (2501; 2502) were amplified with primers listed in Table 1. Each of the receptors was cloned into vector pME18S (kind gift of N. Yamanaka, Japan) through EcoRI and XbaI digestion sites with addition of Kozak sequence, initiating translation, incorporated at the 5’end. pME18S-PTHR1 or pME18S-PTHR2, vector bearing promiscuous G_α15_ subunit (pG_α15_) and plasmid encoding Ca^2+^ reporter protein – aequorin, were transfected into CHO-K1 cell line using FuGene HD Transfection Reagent (Promega, WI, USA). CHO-K1 cells were cultured for 24-48h in Dulbecco’s modified eagle medium nutrient mixture F-12 (DMEM/F12) at 37°C with humidity of 5% CO_2_. The cells were detached and incubated for 3h in 5μM Coelenterazine (Promega, WI, USA). Different concentrations of synthetic neuropeptide iPTH (SGISDQRLAELETLAGLKSLRHRLKGISFPVAYGLVDPNKIamide) or orcokinin OK-B1 (TLDKLSGGEYIRALHRLGamide) (Sigma-Aldrich) were loaded into a white 96-well plate (Sigma-Aldrich, St, Louis, USA) and luminescence emitted after incubation of cells with peptides was monitored with GloMax Multi Detection System (Promega, WI, USA). Experiments were performed in three technical and three biological replicates. Acquired data were analyzed in Excel (Microsoft, Redmond, WA, USA) and GraphPad Prism 11.0.1 (GraphPad Software Inc., Boston).

### In situ hybridization

Whole-mount in situ hybridization (ISH) was performed as described previously (Kim et al., 2006). For each studied stage 5-10 individuals were examined. The dorsal cuticle was removed and whole ticks were fixed at RT in 4% paraformaldehyde. The samples were immediately used or stored in 70% ethanol at -20°C. Template DNA fragment for generation of single-stranded digoxygenin-labeled probe was prepared with primers 2346 and 2347 (Table 1). The probe was synthesized using a specific reverse primer 2347 and PCR Dig Probe Synthesis Kit (Roche, Mannheim, Germany; Table 1). Tissues with hybridized probes were incubated with alkaline phosphatase-labelled antibody against digoxygenin. Development of staining was observed under binocular microscope Leica M205 FA (Leica Microsystems, Germany). Stained tissues were either mounted in 100% glycerol or subjected to immunohistochemical staining (IHC). Preparations were observed using Eclipse 600 fluorescence microscope with Nomarski DIC optics and photographed with Coolpix 990 camera (Nikon, Tokyo, Japan).

### Generation of mouse *I. ricinus* iPTH-specific antisera

The iPTH antigen (CAYGLVDPNKIamide) was commercially synthesized with conjugating keyhole limpet hemocyanin (KLH) to its N-terminal (Sigma-Aldrich, USA). 100μg KLH alone or KLH-iPTH conjugate were mixed with Freund’s adjuvant (Sigma Millipore, USA) in 1:1 ratio and emulsified. Before the first dose, pre-immune blood was collected and then 100μl of the conjugate mixture was inoculated subcutaneously into 4-6 weeks old BALB/c mice. Three 100μg doses of the conjugate were administered and blood was collected by venipuncture of the facial vein two weeks after each dose. Collected antisera were tested on *I. ricinus* tissues with pre-immune serum and KLH serum as negative controls.

### Generation of guinea pig polyclonal sera against *I. ricinus* iPTH

For the initial dose, 100μg of the KLH-iPTH conjugate or KLH alone was mixed with complete Freund’s adjuvant (Sigma Millipore, USA) in a 1:1 ratio and emulsified. Before the initial dose, pre-immune blood was drawn from the lateral saphenous vein. 200μl of the emulsion was administered subcutaneously in 300g female Dunkin-Hartley guinea pigs (n=2/group). For the second and third doses, 50μg of the conjugate or KLH alone was mixed with incomplete Freund’s adjuvant (Sigma Millipore, USA) in a 1:1 ratio and emulsified. 200μl of the vaccinal emulsion was administered subcutaneously. Two weeks following the second and the third dose, immune sera were tested for peptide-specific antibodies by whole-mount immunofluorescence using tissues of *I. ricinus* tick. Pre-immune serum anti-KLH sera were used to screen for non-specific immunoreactivity.

### Wholemount immunohistochemistry

The whole-mount IHC staining was performed as described previously (Medla et al., 2023). Fixed tissue samples were treated with one or more primary antibodies generated from different animals, listed in Table 2. For visualization of bound antibodies, secondary antibodies Alexa-Fluor-488-labelled donkey anti-rabbit IgG, Alexa-Fluor-594-labelled donkey anti-mouse IgG, and Alexa-Fluor-488-labelled goat anti-guinea pig IgG were used. Tissues mounted in 100% glycerol were observed and scanned using confocal microscope system TCS SPE confocal system (Leica Microsystems, Germany) with 488, 532 and 635 nm lasers for excitation. Alignment of identical images from light and fluorescence microscopy and labelling was performed with Adobe Photoshop version 27.6 (Adobe Systems, CA, USA).

**Table 2.**
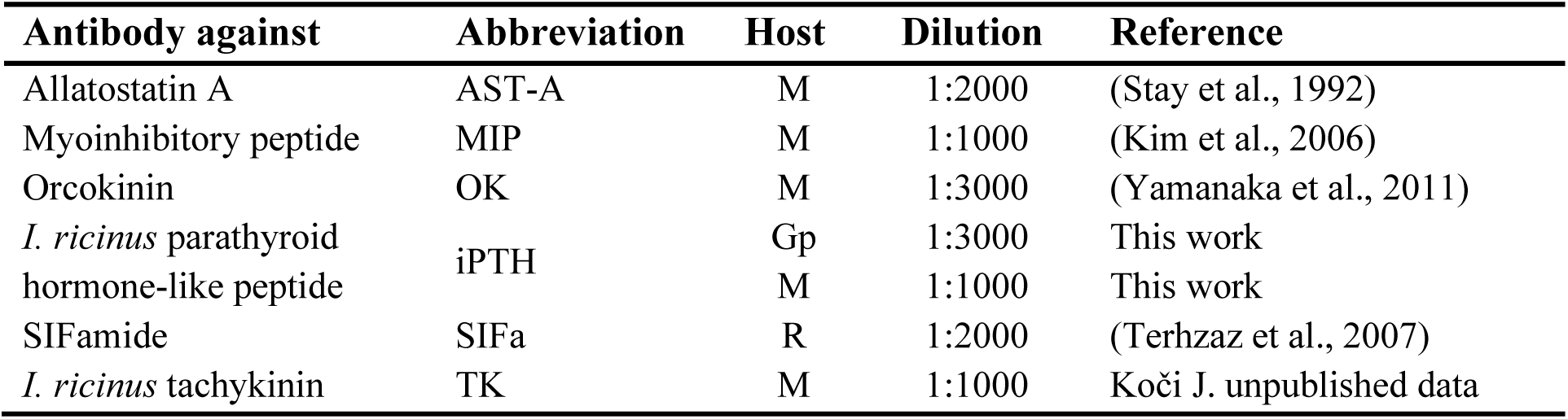
Primary antibodies used in this study. Antibodies were produced in mouse (M), rabbit (Rb) or guinea pig (GP).

### Real-time quantitative PCR (RT-qPCR)

The Real-time quantitative PCR (RT-qPCR) was performed as described previously (Medla et al., 2023). cDNA of different stages and tissues of *I. ricinus* were used (see cDNA cloning and rapid amplification of cDNA ends (RACE)). Each RT-qPCR reaction ran in two technical and two biological replicates. Primers used for RT-qPCR analysis are listed in Table 1. Levels of analyzed transcripts were normalized to reference genes RpS4 and EF1 (Koči et al., 2013; Nijhof et al., 2009). Data were analyzed using 2^-ΔΔCt^ method (Livak & Schmittgen, 2001) and statistical analyses was performed by one-way analysis of variance (ANOVA) followed by Tukeys multiple comparison test in GraphPad Prism version 11.0.1 (Massachusetts, USA).

### RNA interference-mediated knockdown of PTHRs

RNA interference was achieved by administration of gene-specific fragments of double stranded RNA into ticks. Two fragments were designed for each PTHR and a fragment of green fluorescent protein (GFP) was used in a control group. Specific fragments were amplified in PCR reaction with T7 promoter flanked primers. MEGAscript™ T7 Transcription Kit (Invitrogen by Thermo Fisher Scientific, Lithuania) was used to prepare dsRNA and purified with MEGAclear™ Transcription Clean-Up Kit (Invitrogen by Thermo Fisher Scientific, Lithuania). Quality and concentration of the resulting dsRNA was verified by gel electrophoresis and Nanodrop2000 (Thermo Fisher Scientific, MA, USA).

Unfed 1-6 months old *I. ricinus* nymphs were inoculated intra-rectally with dsRNA for GFP or PTHRs. Each nymph was injected with 50nl of dsRNA (3 μg/μl) using Nanoject II (Drummond, PA, USA) and fine glass capillary tip. Nymphs were stored overnight at 22°C and 95% humidity and then administered to the murine hosts. Nymphs (10-12 per mouse) were fed on BALB/c mice until repletion. Three mice were used in GFP group and 6 mice in PTHRs group. Fully engorged released nymphs were stored at 22°C, 95% humidity and 10:14h light:dark cycle. Second dose of dsRNA was administered to replete nymphs 10 days after feeding. RNAi booster was inoculated per coxum in a single 50nl dsRNA dose with concentration of 3 μg/μl. 15 days after the end of feeding, three nymphs per mouse were used for RNA extraction followed by cDNA preparation (see cDNA cloning and RACE). The RNAi-mediated knockdown was verified by RT-qPCR.

### Data visualization

The final figures were prepared using Adobe Creative Cloud software (Adobe Photoshop, Illustrator and InDesign).

## Results

### Identification and characterization of the peptide precursor and its receptors

In this study, we identified iPTH and two putative receptors in the genome of *I. ricinus*. The iPTH transcript gives rise to a 131 amino acid prepropeptide containing a single iPTH (41aa) with a dibasic cleavage site on the N-terminus (RR), and cleavage and amidation sites on the C-terminus (GKR) (Fig. 1).

**Fig. 1.**
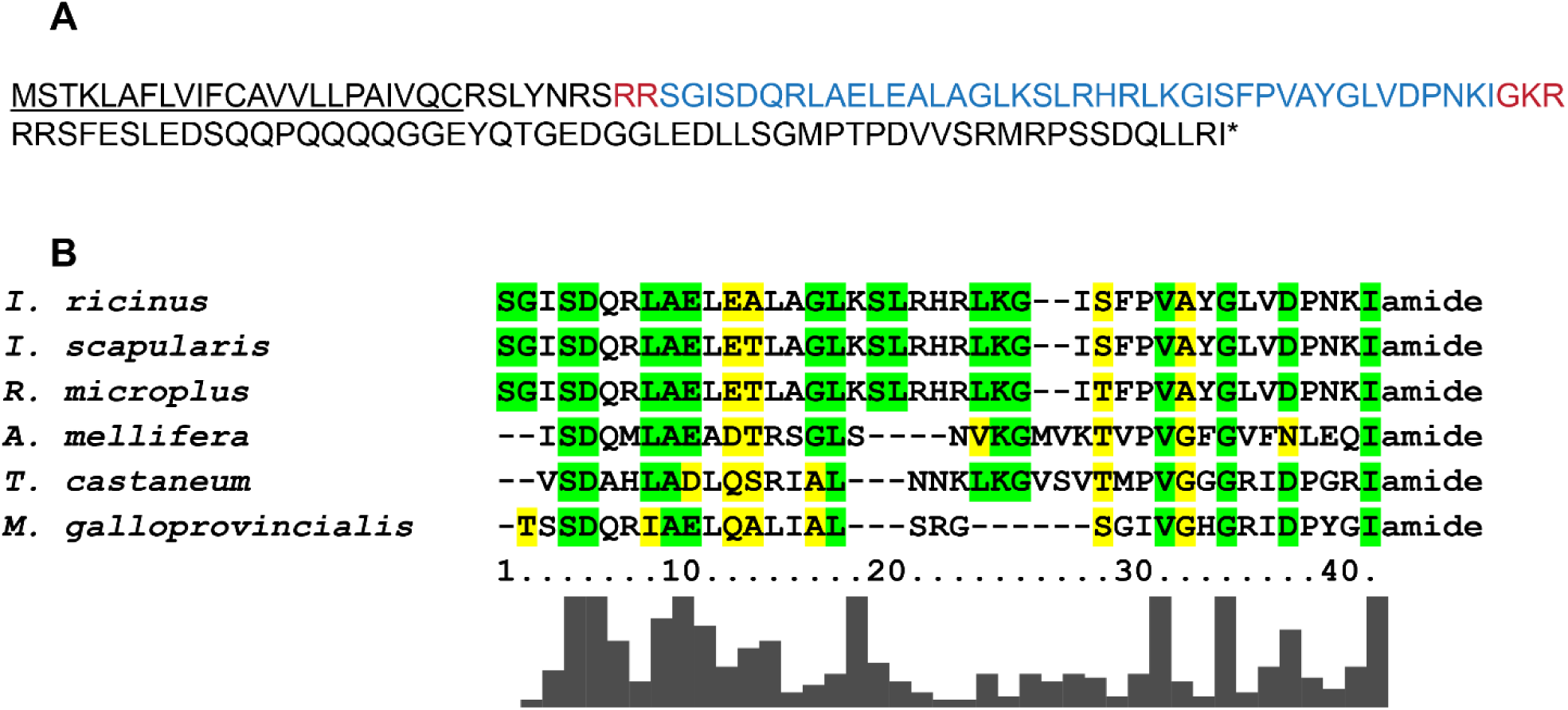
Amino acid sequences of iPTH from ticks, insects and mollusc. A) The sequence of prepropeptide composed of a signal peptide (underlined), cleavage sites marked with red and the sequence of active peptide in blue. B) Sequence comparison of iPTH from the ticks *I. ricinus*, *I. scapularis* (XP_040063819.1), *R. microplus* (UKD60448.1), the beetle *T. castaneum* (XP_008194702.1), the honey bee *Apis mellifera* (XP_026300078.1), and the bivalve mussel *Mytilus galloprovincialis* (VDI22276.1) made with ClustalX 2.1 and green/yellow (identity/similarity) coloring scheme.

Orthologs of mammal PTH receptors were previously found in the beetle *T. castaneum* and other insects (Li et al., 2013b). Based on *T. castaneum* PTHRs we found 5 related putative GPCR sequences (XP_029845012.1; XP_029838024.1; XP_029838018.1; XP_029844994.1; XP_029834640.1) in the genome of *I. scapularis* and two putative receptors for *I. ricinus* iPTH. The proposed *pthr1* transcript includes 1386bp long open reading frame (ORF). PTHR1 is a GPCR of cI28897 superfamily and contains hormone receptor ligand-binding domain smart00008 on the N-terminus (Sup.1). The second GPCR (PTHR2) transcript includes 1656bp ORF. Unlike PTHR1, PTHR2 has a pfam02793 ligand-binding domain on the N-terminus (Sup. 1). To determine specificity of *I. ricinus* PTHRs, we transiently expressed these GPCRs in CHO-K1 cell line for the aequorin-based calcium-mobilization assay. The assay confirmed specific activation of both PTHRs with iPTH in concentration-dependent manner (Fig. 2). Surprisingly, despite the different ligand-binding domains (smart00008 and pfam02793), iPTH had similar affinity to PTHR1 (EC_50_=62,6nM) and PTHR2 (EC_50_= 51,6nM). Neither of the tested receptors responded to an unrelated peptide orcokinin.

**Fig. 2.**
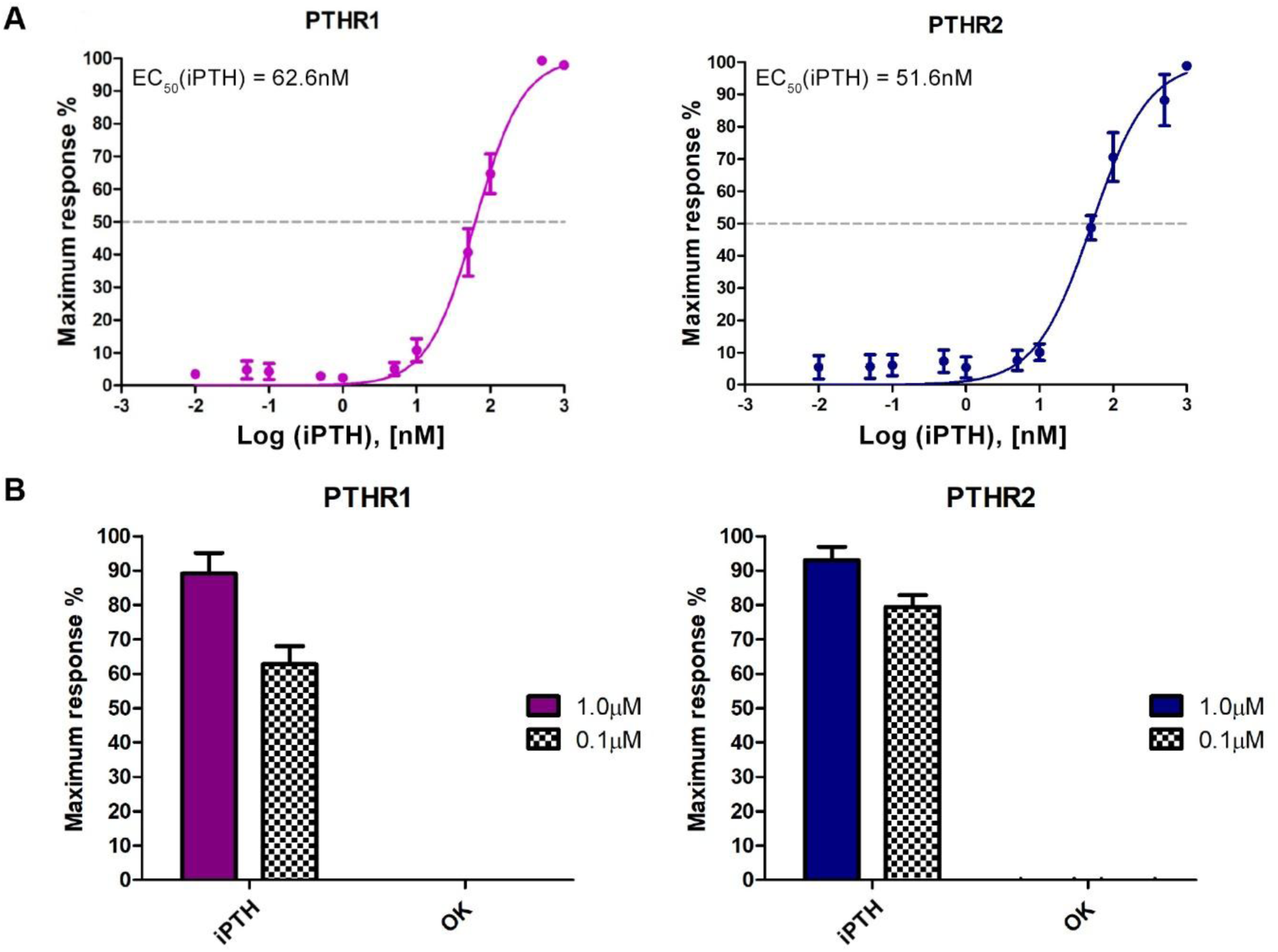
Characterization of iPTH receptors. A) Dose-response curve of PTHR1 (left) and PTHR2 (right) heterologously expressed in CHO cells. Activation of the receptor is presented as a percentage of maximum peak luminescence achieved with the ligand. Each data point is a mean value ±SE (n=3). B) Response of PTHR1 (left) and PTHR2 (right) after application of alternative ligand (OK-B1) normalized against the response to 1μM or 0,1μM of iPTH.

### Expression of iPTH in the synganglion

Utilizing ISH, we detected iPTH transcript in several types of neurons in the synganglion (Figs. 3A,A’). For neuron determination we used nomenclature previously used for the tick *R. appendiculatus* (Šimo et al., 2009). We observed ∼30 pairs of small iPTH expressing neurons in the protocerebral (Pc) lobe and 3 pairs of neurons in the cheliceral lobe (Ch). In the region of olfactory lobe, we detected 2 pairs of ventromedial and 2 pairs of ventrolateral cells. Paired large ventro-lateral neurons were consistently labelled in the pedal lobes (Pd_1-4_VL_1_), plus additional smaller neurons were found on the dorsal and ventral sides. In the dorsal part of the opisthosomal lobe (Os), we observed two pairs of the most prominent iPTH neurons OsSG_1, 2_, and a weakly stained pair of PoHG cells. About 11 other smaller weakly stained neurons were detected in this lobe. We confirmed iPTH production in these neurons with IHC staining (Fig. 3B, B’). Using iPTH-specific antisera we detected axons of neurons innervating the legs (Fig. 3C), chelicerae (Fig. 3D), salivary glands (Fig. 3E) and hindgut (Fig. 3F). Schematic drawing of iPTH-expressing neurons with depicted cells innervating peripheral organs is shown in Fig. 4A.

**Fig. 3.**
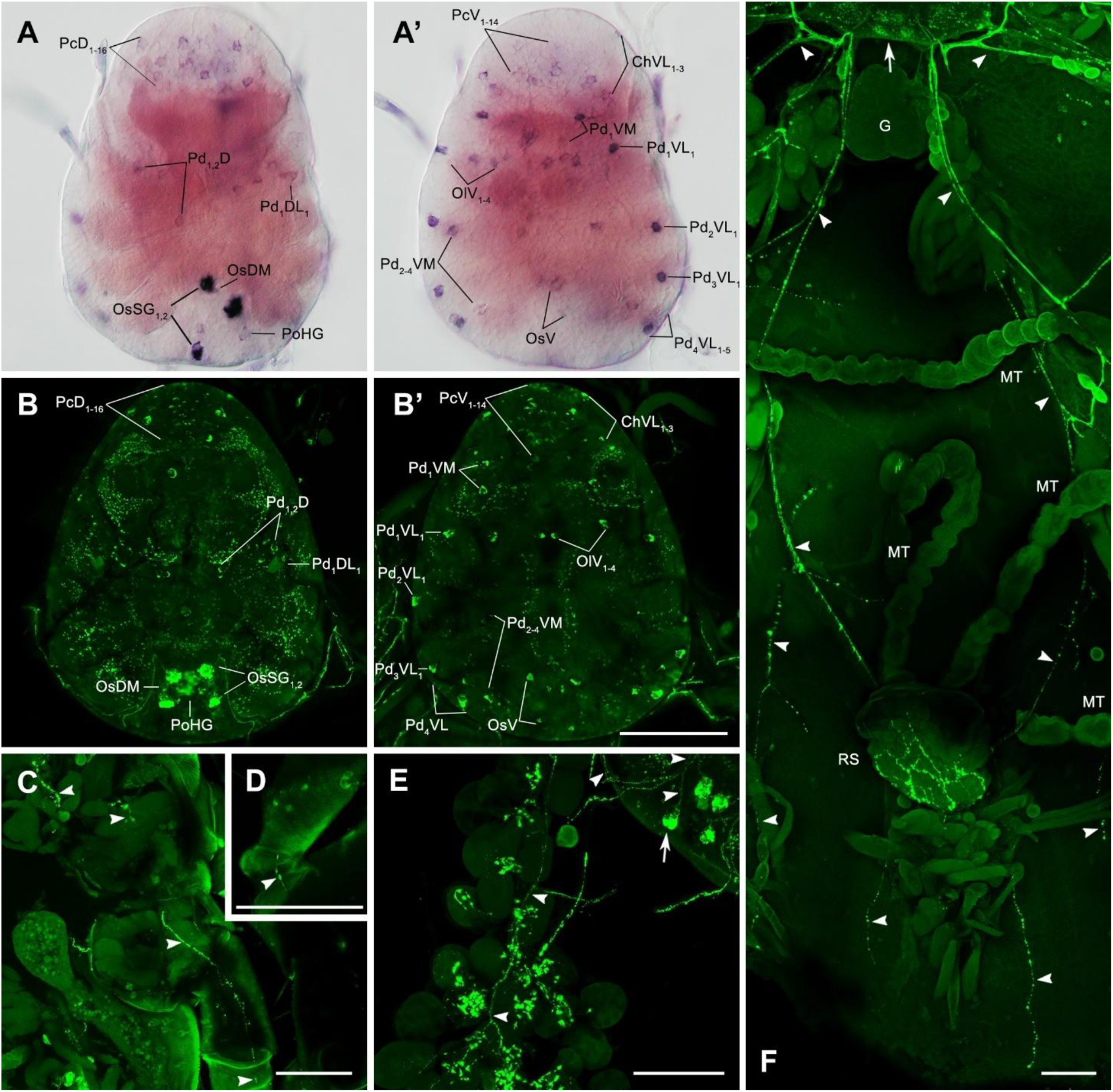
Expression of iPTH in central neurons innervating peripheral organs. iPTH detected by ISH (A, A’) and IHC (B, B’) in multiple neurons of the dorsal (A, B) and ventral (A’, B’) synganglion. IHC staining with iPTH antiserum revealed axons (arrowheads) of iPTH neurons (arrows) that innervate legs (C; 3^rd^ leg); chelicerae (D); salivary glands (E); and complex axonal varicosities on the rectal sack (F). Unfed nymphs (A-E), engorged and detached nymph (F). Malpighian tubules (MT), rectal sack (RS) and gonads (G). Scale bar; 100μm.

**Fig. 4.**
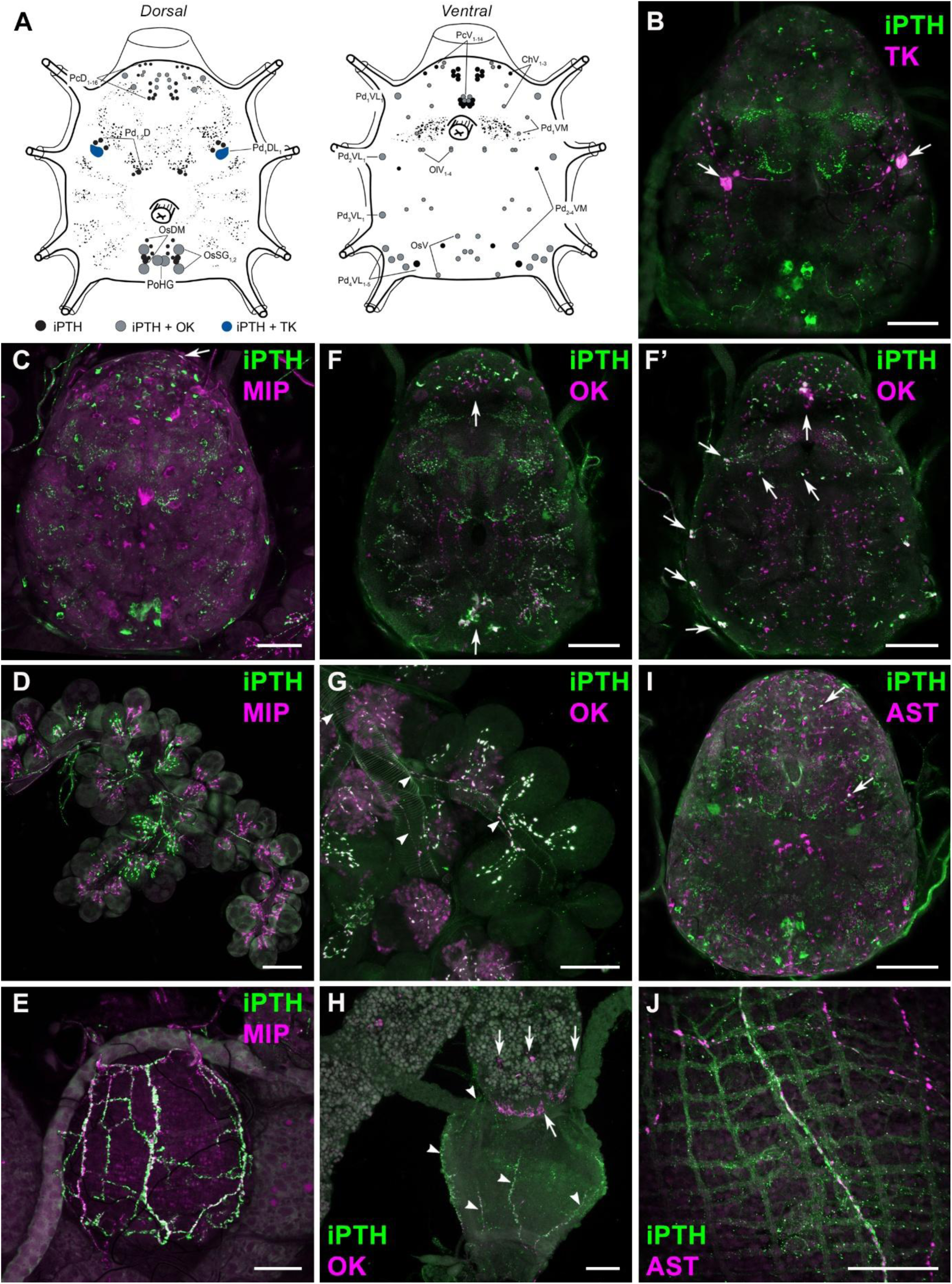
Co-localization of iPTH with other peptides. A) Schematic drawing of *I. ricinus* synganglion with marked iPTH (black), iPTH and TK (blue), and iPTH and OK (gray) expressing cells. B) Co-localization (white) of iPTH (green) and TK (magenta) antibodies in Pd_1_DL_1_ cells of unfed nymph’s synganglion – only *dorsal* section of the brain is shown. Double IHC staining of unfed nymph tissues with antibodies against iPTH (green) and MIP (magenta) is shown in C, D, E. Overlay of the signals (white) was detected only in a few further uncharacterized Pc neurons of the synganglion (arrows in C) and no colocalization was detected in the innervation of salivary glands (D) or rectal sack (E). F, F’) Double IHC staining with antisera against iPTH (green) and OK (magenta) show colocalization of the signals (white) in numerous neurons of the synganglion (arrows) (F dorsal; F’ ventral sections), and in the innervation of salivary glands (G) and rectal sack (arrowheads; H). Note the midgut-rectal sack endocrine cells (arrows). (I) Double IHC staining with antibodies against iPTH (green) and AST-A (magenta) revealed colocalization of signals in only a few small Pc and Ol neurons (arrows). These antibodies did not detect any overlay in innervation of the rectal sack (J). Scale bars: 50μm

### Characterization of neurons expressing iPTH

We used double-IHC staining with various neuropeptide antibodies (Table 2) to identify neurons expressing iPTH. A single pair of large neurons resembled the major lateral neurons producing tachykinin-like neuropeptides (TK) (Šimo et al., 2009; Mateos-Hernández et al., 2021). Double-IHC staining with antibodies against iPTH and TK confirmed signal overlay in these cells and therefore we named them Pd_1_DL_1_ (Fig. 4A,B). Previously, several neuropeptides were identified in the innervation of the salivary glands type II, III (Kim et al., 2018; Roller et al., 2015; Šimo et al., 2009). We detected colocalization of iPTH- and OK-IR in previously identified OsSG_1,2_ neurons that innervate exclusively the acini type II (Figs. 4B,C). iPTH- and OK-IR was also observed in PoHG neurons innervating the hindgut (Figs. 4D,E), while iPTH-IR showed no overlap with AST-A- and MIP-IR (Figs. 4F-J). We further identified enteroendocrine cells showing OK-IR in the lining of the sphincter between midgut and rectal sack (Fig. 4H). Based on their position we named them MRS (midgut-rectal sack) cells. Interestingly, double IHC staining with antibodies against iPTH and OK further revealed signal overlay in numerous neurons of the synganglion (22 Pc, 6 Ch, 6 Ol, 22 Pd and 6 Os neurons) summarized in Fig. 4A.

### Changes of iPTH-IR in innervation of the salivary glands during feeding

We used antibodies against iPTH and SIFa to detect possible release of these peptides from axonal projections innervating the salivary glands during feeding of nymphs and adult females. In nymphs we did not detect obvious waning in the intensity of staining with both antibodies during or after feeding (Figs. 5A-D). Nevertheless, we detected changes in the number of varicosities in the acini of feeding nymphs. Feeding was associated with enlargement of acini and increased length of axonal projections. We also observed a doubling in the number of varicosities within acini II on day 1 of feeding (Fig. 5B), which apparently decreased at the end of feeding on day 3 and in depleted nymphs (Figs. 5C,D). The decrease of SIF-IR varicosities was even more dramatic in feeding and depleted nymphs (Figs. 5A-D). These data suggest iPTH and SIFa release during feeding of nymphs. In adult females the original accumulation of iPTH-IR in unfed and feeding ticks on day 1 remarkably decreased on days 3-5 and only diminutive traces of iPTH-IR were observed at the end of feeding on days 6-7 (Fig. 5G, G’). These data indicate apparent iPTH release during the “slow” phase and its depletion during the “rapid” phase of feeding in adult females.

**Fig. 5.**
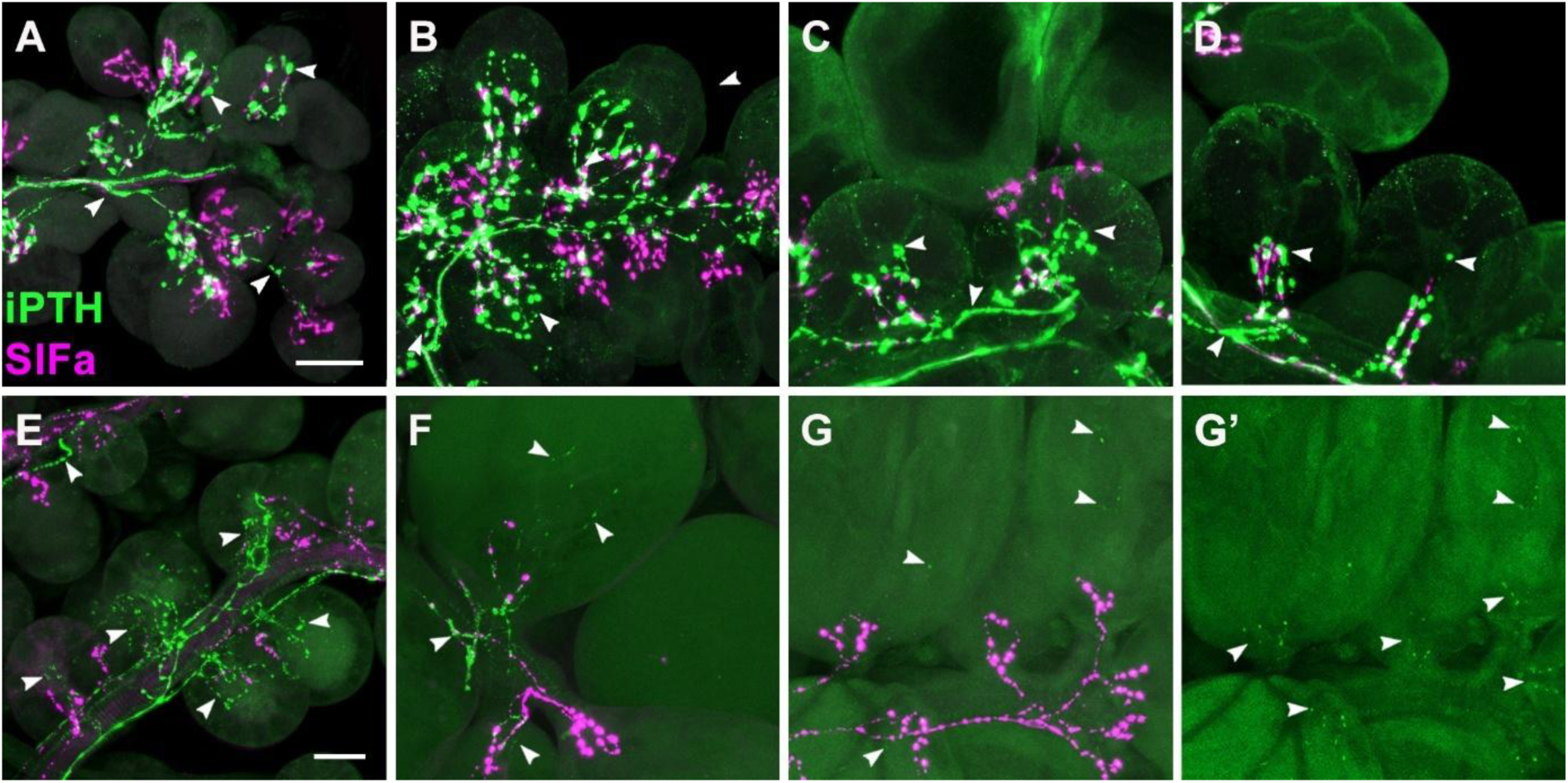
Stage specific differences in immunoreactive axons innervating the salivary glands. Double IHC staining of the salivary glands in nymphs (A-D) and adult females (E-G) with iPTH (green) and SIFa (magenta) antibodies. The intensity of iPTH-IR axons (arrowheads) is prominent in all studied nymphal stages - non-feeding (A), feeding on day 1 (B), feeding on day 3 (C), and replete/detached nymph (D). Strong IHC staining of female salivary glands at the beginning of feeding on day 1 (E), considerably decreased at the end of slow phase on day 5 (F) and was depleted at the rapid phase on day 6 (G, G’). Note very weak/depleted iPTH-IR alone in G’. Scale bar: 25μm

### Expression profile of PTHRs

Using RT-qPCR analysis of female tissues, we detected transcript of iPTH precursor exclusively in the synganglion supporting our findings with ISH (Fig. 6A). Likewise, the highest levels of transcripts of both receptors (PTHR1,2) were detected in the CNS, but increased levels of PTHR2 were also detected in the salivary glands, gut, and female gonads (Fig. 6B, C). We also examined expression of PTHR2 in different parts of the digestive tract, since only the hindgut is innervated by axons producing iPTH. As expected, we found increased PTHR2 expression in the hindgut, which is consistent with iPTH-IR detected by IHC in the innervation of the rectal sac (Fig. 6D). Comparison of PTHR2 expression in the gut of nymphs and adults revealed the highest levels of the transcript in the repleted and detached nymphs (Fig. 6E). Detailed examination of PTHR2 expression in the gut of nymphs showed only low levels during feeding, which considerably increased after repletion and detachment on days 7 and 14 (Fig. 6F). Further developmental analysis revealed high levels of PTHR2 expression in the salivary glands of feeding females, which declined after detachment from the host (Fig. 6G). Interestingly, in the female gonads PTHR2 expression increased from the background levels at the beginning of feeding to very high levels at the end of blood intake, then considerably declined after detachment and increased again during egg laying (Fig. 6H).

**Fig. 6.**
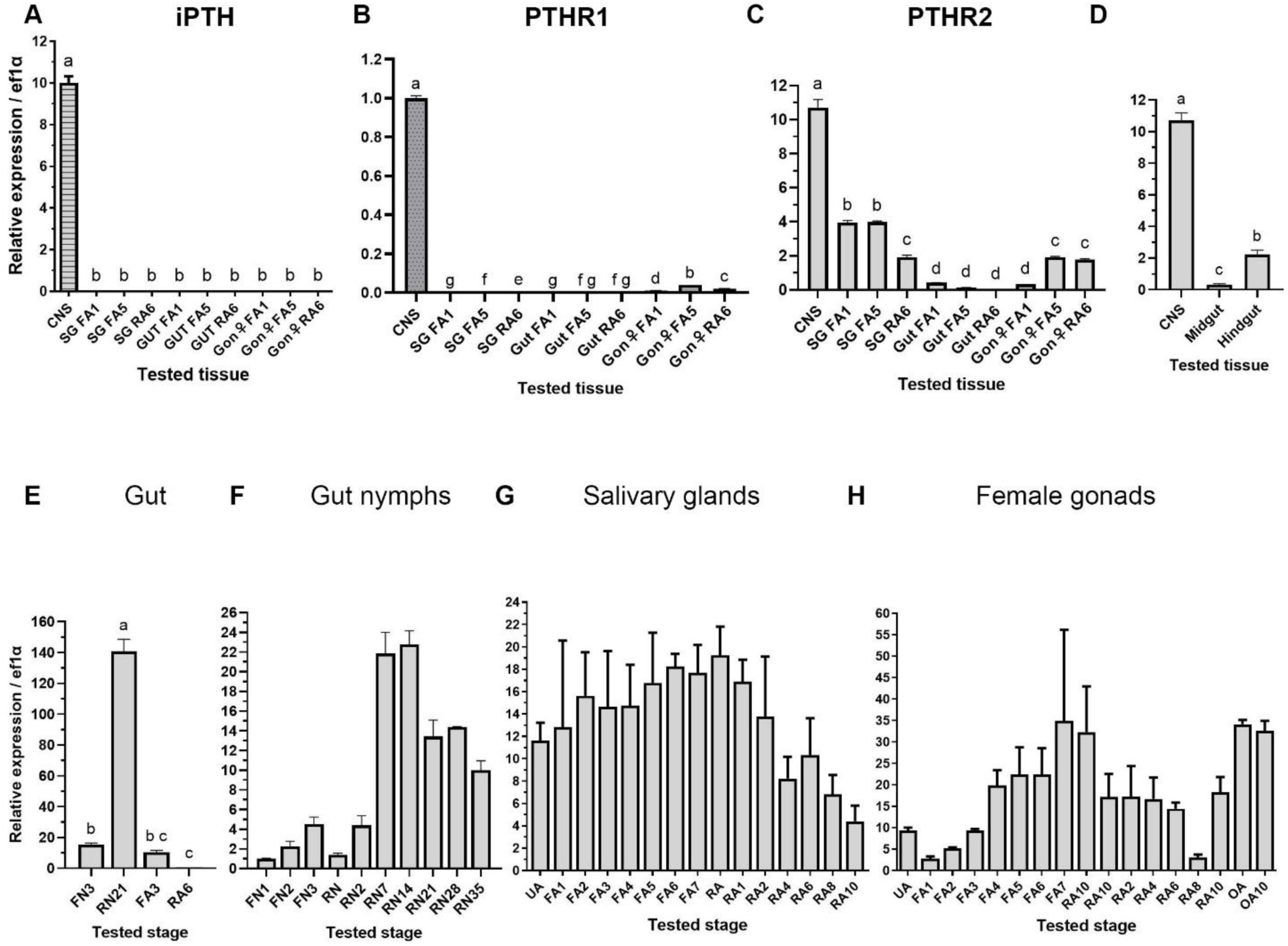
Expression of iPTH and PTHRs in nymphs and adult female. Differential expression of iPTH (A), PTHR1 (B) and PTHR2 (C) in the synganglion, salivary glands (SG), gut, and gonads (Gon ♀) of female *I. ricinus*. Normalized tissue specific expression of 1 day (FA1), 5 days of feeding (FA5), and female fed to repletion 6 days post-detachment (RA6), is compared to expression of the respective genes in mixed sample of synganglia of all stages. D) Comparison of PTHR2 expression in the CNS with expression in the midgut (MG) and hindgut (HG) of unfed female. E) Levels of PTHR2 in the gut of 3 days feeding nymphs (FN3), replete nymphs 21 days post-detachment (RN21), 3 days feeding female (FA3) and replete female 6 days post-detachment (RA6) are shown. F) Stage specific expression of PTHR2 in the gut of 1-3 days feeding nymphs (FN1-3), replete nymphs on days 0, 2, 7, 14, 21, 28, and 35 days post-detachment (RN0-35). Stage-specific expression of PTHR2 in salivary glands (G), and female gonads (H) – unfed (UA), 1-7 days feeding (FA1-7), replete females on days 0, 1, 2, 4, 6, 8 and 10 days after detachment from host (RA0-10), female oviposition on day 1 and 10 (OA, OA10). Error bars indicate standard error of the mean (n = 3). The data were analyzed with one-way ANOVA with Tukey’s multiple comparison post hoc test and visualized with Compact Letter Display (p value <0.0001).

### RNA interference-mediated knockdown of PTH receptors

To determine possible role(s) of *I. ricinus* PTHRs, we utilized RNA interference in nymphs and observed phenotypic changes in adult ecdysis as a result of PTHRs knockdown (KD; Fig. 7A). Unfed *I. ricinus* nymphs were injected with PTHRs dsRNA or GFP dsRNA (control group) and were allowed to feed on murine host till repletion. Although the dsPTHRs group fed significantly slower than the control group (Fig.7B), weights of replete nymphs were not altered (Fig.7C). To ensure PTHRs KD over a long period of time, ten days post-feeding the nymphs were inoculated with the second dsRNA dose. To evaluate KD of PTHRs, mRNA levels were tested from the whole bodies. The efficiency of PTHRs KD varied individually but expression of PTHR1 was reduced by 43-94 %, and PTHR2 by 41-80 % compared to control (Fig.7D).

**Fig. 7.**
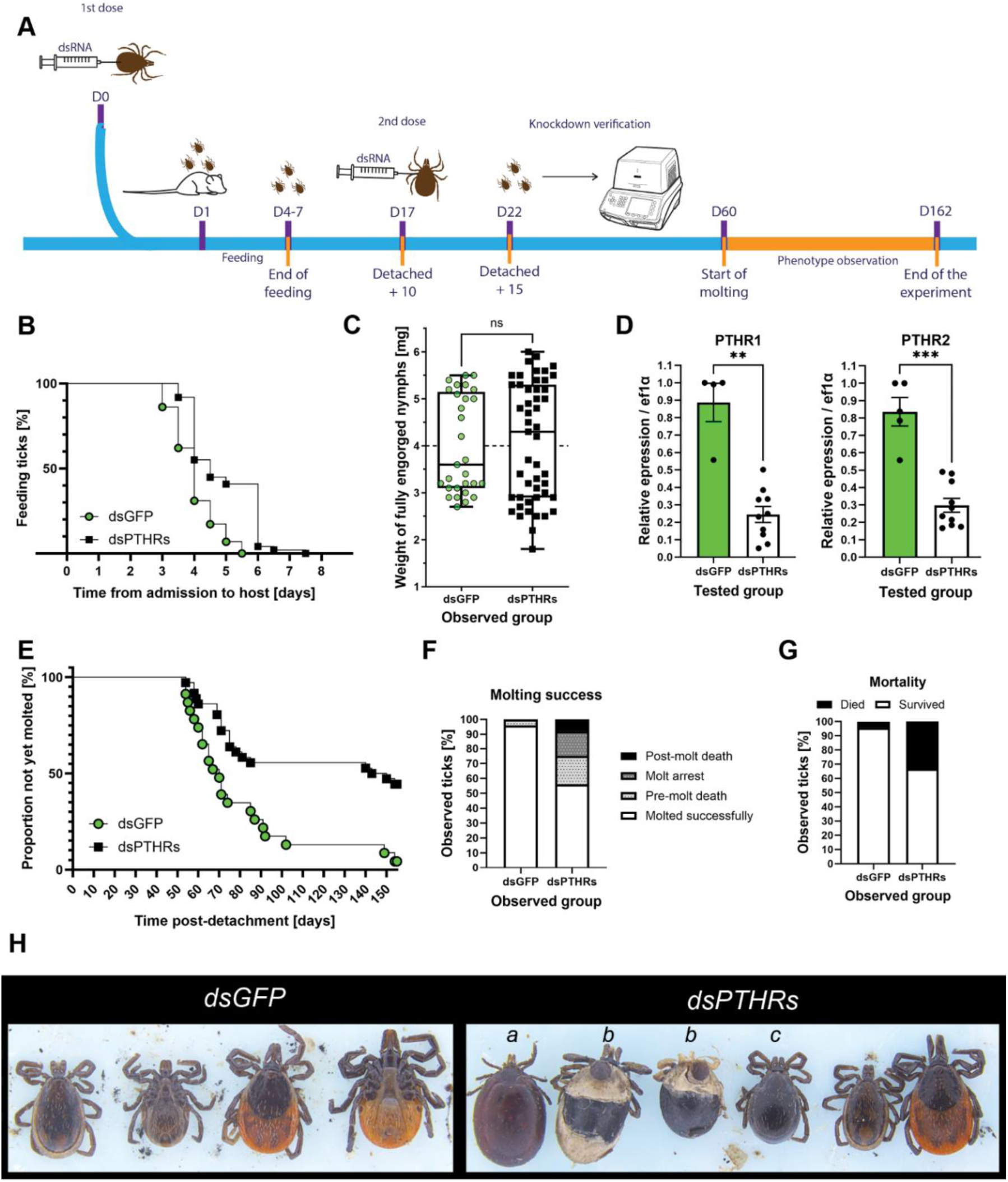
RNA interference of PTHRs in nymphs of *Ixodes ricinus*. The workflow of RNAi PTHRs is summarized in A. B) Nymphal feeding on murine host was observed with Kaplan-Meyer curve with Mantel-Cox Logrank test (χ2 = 19.46; p < 0.0001; ****). C) Weight of fully engorged nymphs was analyzed using t-test with Welch’s correction (p value 0.7279; ns). D) Verification of PTHRs RNAi mediated knockdown was tested on five (dsGFP) and ten (dsPTHRs) randomly selected replete nymphs, five days after the second dsRNA dose (15 days after the end of feeding). Data were analyzed with Mann-Whitney test PTHR1 p = 0.0020 **; PTHR2 p = 0.0007 ***. Error bars indicate Mean with SEM. E) Molting of the fully engorged nymphs was observed over 5 months (155 days) after repletion. The molting is presented with Kaplan-Meyer curve with Mantel-Cox Logrank test (χ2 = 13.98; p = 0.0002; ***). F) The molting success in PTHRs group compared to dsGFP control. Data were analyzed with Chi-square test (χ2 = 45.31; p < 0.0001; ****) and Fischer’s exact test (p < 0.0001; ****). G) Overall survival in dsPTHRs group compared to dsGFP control group. Mortality was evaluated with Fischer’s exact test (p = 0.0051; **). H) RNAi phenotypes of the ticks – pre-molt death (a); molt arrest (b); post-molt death (c).

The PTHRs knockdown (KD) caused mortality in three different stages of ecdysis; only 63.89% ticks completed successful ecdysis compared to 95.65% in the control group (Fig. 7E) and remaining ticks died. The mortality of KD ticks increased even before ecdysis; 19.44% in dsPTHRs group and 4.35% in dsGFP group. 16.67% of dsPTHRs nymphs entered molt arrest before ecdysis initiation, while no arrest was observed in dsGFP group and 8.33% of dsPTHRs adults died after successful ecdysis compared to 0% in dsGFP (Fig. 7F). In conclusion, in all experimental ticks (nymphs and adults) we observed higher mortality in dsPTHRs group (44.44%) compared to control group (4.34%) (Fig. 7G).

## Discussion

As in other arthropods, tick life cycle comprises multiple molting events that are tightly regulated by coordinated hormonal and gene expression changes. Based on our data, we propose that iPTH signaling plays a role in ticks’ molting. RNAi-mediated knockdown of iPTH receptors in nymphs resulted in prolonged feeding, increased mortality and ecdysial arrest, supporting the involvement of this pathway in the molting process. Consistent with findings in *T. castaneum*, this phenotype may be linked to disruptions in chitin metabolism (Xie et al., 2020). Beyond its proposed role in molting, our data suggest that iPTH signaling has broader physiological functions in ticks.

We identified a single iPTH in *I. ricinus* (Fig.1), which shows high sequence conservation with previously identified iPTH in *R. microplus* (Waldman et al., 2022). In addition, we identified and functionally characterized two PTH receptors in *I. ricinus* that show high degree of homology to *T. castaneum* iPTHRs (Supl Fig. 1) (Li et al., 2013). Our data indicate that *I. ricinus* iPTH shows similar affinity to both receptors (Fig. 2) and exerts its functions by specific activation of iPTHR1 in the CNS or iPTHR2 in the CNS, SG, hindgut and gonads (Figs. 6B-G). This contrasts with the finding in *T. castaneum* in which iPTH preferentially activated PTHR1 predominantly expressed in gut and showed reduced affinity to PTHR2 predominantly expressed in CNS of nymphs or gut of adults (Xie et al., 2020). Contrary to the expression profile of iPTH in *T. castaneum* or the distribution of *I. ricinus* PTHRs, RT-qPCR showed iPTH expression only in tick’s synganglion (Fig. 6A) suggesting its neuroendocrine mode of action (Xie et al., 2020). Based on receptor distribution, iPTH is probably involved in multiple physiological processes associated with feeding and reproduction. Furthermore, detection of iPTH-IR axons innervating distant parts of the tick body including muscles of chelicerae and legs (Fig. 3) indicates its role in locomotion and movements of external organs.

Colocalization of iPTH with TK in Pd_1_DL_1_ neurons, which arborize on the neurohaemal periganglionic sheat (Fig.4B) suggests that a cocktail of these peptides is released into the hemolymph to target peripheral organs. In insects, TKs are pleiotropic neuropeptides involved in regulation of lipid metabolism, diuresis, locomotion, olfactory sensing, in male aggressive behavior (Bubak et al., 2019; Gui et al., 2017; Skaer et al., 2002; Song et al., 2014). The function of TKs in ticks remains unknown we speculate that they may be involved in similar processes (Mateos-Hernández et al., 2021).

Of particular interest is colocalization of iPTH with OK-IR in multiple interneurons and neurosecretory cells of the synganglion. In insects, OKs have been implicated in the regulation of the prothoracic gland, cuticle pigmentation and oogenesis, thereby contributing to insect development (Ons et al., 2015; Xie et al., 2020; Yamanaka et al., 2011; Wang et al., 2019), which is consistent with our RNAi-derived molting phenotype. Furthermore, OKs are known as “awakening” peptides; and function in regulation of circadian rhythms, or cuticle pigmentation, etc. (Hofer & Homberg, 2006; Jiang et al., 2015; Ons et al., 2015; Wang et al., 2019). Colocalization of iPTH and OK-IR in OlVM_1_ and OlVL_1,2_ (Fig.4A, F’) suggests a potential role of these peptides in olfaction. These peptides are further colocalized with AST in a pair of OlVL_1_ neurons (Fig.4I), which in insects plays role in appetitive olfactory learning (Urlacher et al., 2016; Yamagata et al., 2016), further supporting the possibility that these neurons contribute to sensory-behavorial intergration in ticks. Ticks’ olfactory lobe processes information from Haller’s organ and thus allow ticks to interact through semiochemicals and react to external stimuli (Menezes et al., 2021). Consistently with previous results on OK (Roller et al., 2015), iPTH/OK-IR was detected in neurosecretory cells OsSG_1,2_ of the opisthosomal lobe which innervate salivary glands (Fig.3, 4). Neuropeptidergic control of these organs has been intensively studied (Kim et al., 2018; Roller et al., 2015; Šimo et al., 2009, 2012; Šimo & Park, 2014; Vancová et al., 2019). We show that iPTH/OK-IR (and PDF-IR) of the salivary glands originate in OsSG_1,2_ and axons of these neurons innervate acini type II (Fig.3-5). The salivary glands control a variety of functions crucial for ticks’ physiology. Acini type II contain 6 types of granular cells secreting diverse products associated with e. g., cement production or evading host’s immune system (Binnington, 1978; Coons & Roshdy, 1973). These granular acini change shape and size during feeding (Binnington, 1978). Interestingly, according to previous study PDF-IR in acini type II are in direct contact with myoepithelial cells and abluminal epithelial cells suggesting their control by these axons during feeding (Vancová et al., 2019). According to our IHC staining, iPTH is released from axonal projections in the beginning of feeding in females (Fig.5E-H) suggesting it’s importance in the initial phases of adults’ feeding. Similar pattern was detected for OK-IR in *I. ricinus* female (Roller et al., 2015).

iPTH-expressing neurons of opisthosomal lobe project their axons into the muscles of rectal sack (Fig.3). Double IHC staining with various antibodies proved these cells to be PoHG neurons expressing iPTH and showing OK-IR (Fig.4) (Roller et al., 2015). This connects iPTH to one of the 4 known types of neuropeptide innervations of rectal sack – 1. originates in PoHG (iPTH/OK-IR); 2. originates in PoHG_1,2_ (MIP/SIFa-IR); 3. originates in OsHG neurons (FGLa/AST-IR); and the 4. innervation is provided locally from peripheral neurons at the base of rectal sack (sNPF-IR) (Šimo & Park, 2014; Roller et al., 2015; Medla et al., 2023). Hindgut and Malpighian tubules are major excretory organs in tick’s body (Sonenshine, 2014). It has been shown that contractions of hindgut are under control of MIP and SIFa from PoHG_1,2_ (Šimo & Park, 2014). In *Orconectes limosus*, OK directly affects spontaneous contractions of the crayfish’s hindgut (Stangier et al., 1992). Based on these observations, the newly identified MRS-OK-IR endocrine cells (Fig.4H) and previous reports on OK signaling, we speculate that OK and iPTH may contribute to the regulation of excretion processes potentially via modulation of rectal sack function.

We identified many iPTH-IR axons innervating distant parts of tick’s body including chelicerae, legs and axonal projections arborizing on the cuticular surface in posteroventral region of tick’s body (Fig.3C, D, F). Previously, OK-IR of muscles in the posteroventral region was described (Roller et al., 2015). Taken together, based on site of iPTH expression, its colocalization with other neuropeptides; widespread projections throughout tick’s body; expression patterns of PTHRs; and RNAi-mediated phenotype, we propose iPTH to be multifunctional neuropeptide involved in tick’s development.

## Acknowledgements

The funding for the genome data of *I. ricinus* used in this study were provided by The Autonomous Province of Trento under the EU FP7 People Programme, Marie Curie actions - cofound, postdoctoral project Genotick to Dr. Giovanna Carpi. Special thanks to Drs. M. Derdáková, L. Klučár and M. Stano for making the genomic database of *I. ricinus* accessible for the research. Declaration of generative AI and AI-assisted technologies in the manuscript preparation process. All the authors declare no generative AI was used during the manuscript preparation.

## Funding

This work was supported by the Slovak grant agencies: APVV-21-0431, APVV-23-0168, VEGA-2/0037/23, VEGA 2/0040/23, Slovak Republic Recovery and Resilience Plan No. 09I01-03-V04-00089, TUBITAK-2024-01, and PostdokGrant APD0194. Matej Medla is a recipient of Štefan Schwartz support fund 2025/OV2/018.

## Notes

### Competing Interest Statement

The authors have declared no competing interest.

