## Supplement 1 for "Identification and characterization of iPTH and two parathyroid hormone receptors-like (PTHR1 and PTHR2) in the tick *Ixodes ricinus*"

*Ixodes ricinus* PTHR1 nucleotide sequence

AGTGGACTGGTGGCGGTGTGGCCGGCGGTGCCTGTTGCTCCCTTGGCATCGGGACTCGGCTGGACGTC
GTCGTCGGATATCTTCTCTCTCACTCGTGCGGATCGCTCGGACGTTTCTTGGGGTCGCTTCGAAGACG
TCGCCGCACGCTCTTCGCACGCTGAGTTTTGCTGAACCTTCGCTTCGGTACCTCTTGCTGTCAGCGTC
CCAAGAAAATCCAGCTGGGCCAATCCGATCTAGCCGCCATTTTCG**ATGCAGACGCTCAGTGGCTGGA**
**GCGAGCCAAGCGCGACTGCCGGGTCAACGAGAGCTTCACCGTCGACATTGGATTGAGCGACGAGTCCA**
**GGTTGAACCTGCACGACATAGAGCTCAACTCCACGGGCGGTGACTTCTGTCCGCGGGTCTGGGACGGC**
**ATTCTTTGCTGGGAGAGTGCACCGGCCGACACGACCGCACGCGCGCCATGTCCTGAATACCTCGACGG**
**CTTCGACACCACCCGGACTGCGAGCAAGCTGTGCACGGAGAACGGCACCTGGTACCTCAGCCCTCTGC**
**ATGGCCAGACGTGGACCAACTACTCGCAGTGTTTCCTGGCAAAGACATCGGACAGCCTCTCCATGTTC**
**GAGCCTCACCTCCCGACCATCAAGCTTATTTCGAAGATTGGCTACACAGTGTCCTTCGTCAGTCTCAT**
**TGTCGCTTTCATCATCCTGGCTTCTGTCAAGAAGCTGCGGTGCCCTCGGAACAGCTTGCACATGCACC**
**CGTTCCTGTCTTTCATCTTGCGCGCCCTTGCAGTCTTGGTCAAGGACGCCATCTTTGTGGACGGCATT**
**GGCTTCAGCCCCAACATGGACTTCAGCsCAGGACAACGTGGACTGCAAGGTGTTCACGAGCTTCTGGC**
**ACTACGTGCTGATGGCCAACTACTGCTGGATCCTCATGGAGGGACTGTACCTACACAGTCTCGTCTTC**
**CTCGCCTTCTTCACGTTGGACAGTTCGGGGATCATACGATACGTCGCTCTTGGTTGGGGGCTTCCCAT**
**ACTGTTCCTTGTTCCTTGGGTAGCAGTAAGAGCAATGCTTGAAGACACGTATTGCTGGACAACAAATG**
**ACACAAAATCTTACTTCTGGATTATCCGAGGACCTATCACGGCGTCCATTGTGGTGAACTTCGTCCTA**
**TTTATCAACATTACAAGGGTACTCTTTGTAAAGCTATTCACCACCCAAACCCACCAAGCAAGGAAGTA**
**CAGATACAGGAAGTGGTTCAAGTCGACTCTTGTGCTGGTCCCTTTGTTCGGCGCCCACTACGCCTTTC**
**TCCTCGGAATGTCATTGGCGGCAGCGGGCGACCTTCTTGAGCTGGTGTGGCTCTACATCGACCAGCTG**
**TTCTCTTCATTTCAGGGATTTCTGGTGGCCACGCTCTACTGCTTCCTGAACGCCGAAGTGGGTGCCGA**
**AGCACGTGCCGCCATGGCGAGGCATCGCCGGACGTGCTGCTCGGACGGACTTCCGCTGCGGGTGAACA**
**AGGCTCAGGGCAGTGGTTGCAACGAGCCCAGGGCGCGTCTATCACTGCTCACGCGCTCGCTCACGCTT**
**ATCACCAGGCTAGAGGAGCGGCCACTATCTGCCAAGGAAGACTTCGATATACCACTCCACCAGGTCAC**
**ATAG**CCTGCGAGACGACACCACCTTAACCTCATCGATGAAACGGTGCTCAAACAAATATCCCGACTTT
CCCCCACCTGGATTTTAGGCAAGGCATCAGCACATGAGAACTGACCTTTTCGGCAAACCTGGGCTGGG
AACTCAAGGCTACCTACTCAAGTTGTCGATACCTGACTCGTAATTTGTGCGAATAGTTTTCTGCCGTG
ACAGCTGCGCTTGCCTCCTATCCAAGGAAGTTATTTTAAAGTGTAGCACCCAAGTCTTGGGATTCTGT
TTTTTAATGCATGGAAAACGTTTCTAGCTGAAGAATAAGAAGTGTAATACCATTTACGGGGATAAAAT
TTGTAAAATGTTAAGCACATGTGTACTGGAAAGGTTGCTGTAAACMCATGTTGCACTTTTTGTAGATT
AGTGAAATCTCAAAATCTGTAACGGCACGCT

Aminoacid sequence of the *I. ricinus* PTHR1

MQTLSGLERAKRDCRVNESFTVDIGLSDESRLNLHDIELNSTGGDFCPRVWDGILCWESAPADTTARA
PCPEYLDGFDTTRTASKLCTENGTWYLSPLHGQTWTNYSQCFLAKTSDSLSMFEPHLPTIKLISKIGY
TVSFVSLIVAFIILASVKKLRCPRNSLHMHPFLSFILRALAVLVKDAIFVDGIGFSPNMDFSQDNVDC
KVFTSFWHYVLMANYCWILMEGLYLHSLVFLAFFTLDSSGIIRYVALGWGLPILFLVPWVAVRAMLED
TYCWTTNDTKSYFWIIRGPITASIVVNFVLFINITRVLFVKLFTTQTHQARKYRYRKWFKSTLVLVPL
FGAHYAFLLGMSLAAAGDLLELVWLYIDQLFSSFQGFLVATLYCFLNAEVGAEARAAMARHRRTCCSD
GLPLRVNKAQGSGCNEPRARLSLLTRSLTLITRLEERPLSAKEDFDIPLHQVT*

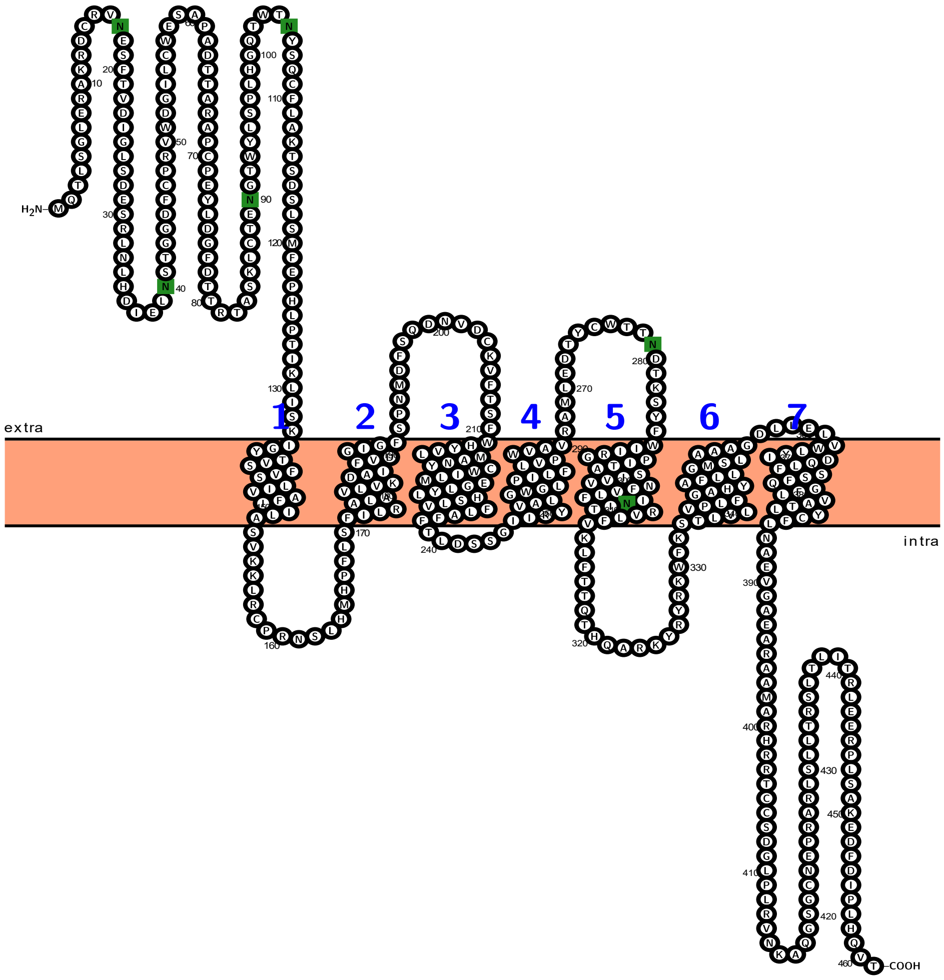


**Fig.1** Graphical visualization of *I. ricinus* PTHR1 made by Protter 1.0. Predicted N-glycosylation motif is marked green. Each of 7 transmembrane helices is numbered in blue.

*Ixodes ricinus* PTHR2 nucleotide sequence

GCGACCGCGTCTGGATAGAGAACCGTGCCTCGCCGGTGGCCTCGTCCGTGTGTTGTGTGCCAGCGTTG
ACTGTGTAGCGACGGCAAAGGAAGAAG**ATGCCCTCCGTCTTCGAGCTTGTCCACAAGAGCGCTCCGCT**
**TCATATTATCTTGGCCGATCAAGACTCCAACATGCAGCTCAGCGCTCTTCAGGAGGCCATGTTGGATT**
**GCCTCAAGAATGACAGCAGCGCAGTTTTTGACTCTGGCGGACTCCCAACATGTCCCGGCGTCTGGGAC**
**GGAATCTTATGCTGGCAACCCGCTCCTGCAGGGACCATGGCCCAGGTCAACTGCCCGGGCTACGTCTA**
**CGGCTTCGACAAACAAGAAATGGCCACTAAGTGGTGCACGGAGAACGGGACATGGTACGTGAGTCCAG**
**AGCACGGTCAGCCCTGGACCAACTACACTCGCTGCTTCCCGCAGGGCGCGACCATGCCCTCCGAGAAT**
**ATCTCCATCTTCGAGCCCCACCTGCCGACCATCAAGCTCATTTCCAAGGTCTGCTACACAGTGTCCCT**
**GGTGACGCTCCTAGCCGCTTTTCTCATGCTGGCTTCCATGAAGAGACTCCGCTGTCCCCGGAACAGCC**
**TACACATGCATTTGTTCATTTCCTTCATCCTGAGGGCGCTAGCATTCCTGCTTAAGGATGCACTCTTT**
**GTCGACGGCATCGGACTTTCGACTAACGTGGACTTCGACAAAGAGAACGTTGACTGCAAGGTGTTCAC**
**GAGCTTCTGGCACTACGTGCTGATGGCCAACTACTGCTGGATCCTCATGGAGGGACTGTACCTGCACA**
**GTCTCGTCTTCCTCGCCTTCTTCACGGACAGTTCGGGGATTATACGATACGTGGCACTCGGATGGGGC**
**CTCCCCGTTTTGTTCATCATTCCCTGGGTCGTGGTGCGAGCGACGCTGGAGGACACCCTTTGCTGGAC**
**GACCAACGTCAGGCAAGACTTGTTCTGGATCATCCGAGGACCAATCACAGCGTCCATCGTGGTGAACT**
**TCGTTCTCTTTCTTAACATCACGCGGGTTCTGTTCGCTAAACTCTTCGCTTCACAGACGCCTCAAGCA**
**CGCAAGTACCGCTACAGGAAATGGTTCAAGTCGACGCTGGTTCTTGTCCCATTGTTCGGGGCGCACTA**
**TGCTTTCCCGCTCGGAATGTCCCCGGCTGCCGCCGGGGACCTCGTGGAGTTGATATGGCTCTACGTTG**

**ACCAGCTCTTCTCTTCATCTCAGGGGTTCGTGGTGGCGCTGCTCTACTGCTTTTTGAACGGCGAGGTG**
**CAGACGGAGCTGCGCAAGTTGTTCCAGAGATGGCGCTGCGACCGTCGCGGTCCGGGCTGCGGGGACAA**
**GCAGCTGCAGCACCGGCACTCGCTCTTCACCCAGTCGGTCACTTTCCTCAGCAGAGGTCGGAGTTCCA**
**TCCAATCTTTGCACTCCGTGGACAGGAGAGAAGGCGTAGCCAAGAGCCCGTCTCCAAACTCCATCAAG**
**AGTAAGATGAGCCAGCGGCTGCTGAGCAACGCCGACCAGGAGTCGCTGGGTGCGGTCAAGAGGATGTC**
**CATTCCGGAGTACCAGGACCCAGCCAAGCGGACGCCCGAGGACAAGACAGGCAGCCAGCGAAAACTCA**
**ACGGAGGCTGTCCGATCCGCTGGGAGTGCCACGATGCAGACTCGGACGGCGGTGCACCAGCCGACACG**
**AGTGAAGACAACCTTCTCGAAGGGGTCATCCAGAAAGAGACGTGCCTCTAA**GGTTAGGCCTTTTGTCG
TGGCTTTCACGGACCGTGTACAGTGATCCCCAGAGGACTGACTGAACCACGCCACTGCCCCCTTGTTG
ATGGAAACGTCGAATTAGCCTACGTTCAGTGATCTCGGTAACCACGTCTGGATAATCATGACCGCTAA
GTGGTTGTGAAGAACTGTGACCAACCAAAAAAGTCAGTAAAATTGGCTGTCTAATTAATTGGGGTTGG
GAACCGGTGGAACCGACTCGACTACTGAGAAAAAATATATCAAACCCTCGATATGAGTACGCAAGAAT
GGCATGTTAAACTTTTACCGTTTATGGAACCATGGTGGAAATATTTTGCCTCAAGAGTTAATCGCATA
GTAACATCATCTGCGAGCATTTTCACCGCCAACCCAGAGCTAATTCGCTTTTAAGAAAGGTGAACGGC
CTGGGACAAACCAACCACCTCTGAACTTTGAAAGTTGACAAAGCGGAATGTGGATTTAGCGGAAACCG
AGTCCAATTTCTCGGGCTATTTATCTGCGGTGAGAAAGTTCCACGATCTCCACATCTTCCTGCGAATA
TCAACGAGGCAATGCTTGAATACCATCAATTGTTGCCACCACACAGTTTTAAGCTCTAACTTGACTGA
AATCGAATACACAGAGTCGTCAATATATTATAAAATGTCTGCCATTCCATTACAAGAAATTATTATTT
CCGCTGACGTTTCTTATTTTTAAAAAACGCCGCCAGCTTAATCATTAACTGGCGATTTCCTACACAGA
GCAGTGGTGAATCCTTTTATTTCTCTATTCGTTTATTTCACGTACCAGTGTCAACCAATTCAAAGCAA
TGGGCGTTCGTGTTCTCGAGAGCGTCAATCGCCAAATAATTATATGCACTCGTACATTTTACTAAATA
TTAACTGTGTTAGGAAGGTATCTCTAAATGTGTTAATGATGATGTTTCCTTGTTGTTAGACACGTGCA
AAATGGATTTTAAACCGGGCTCAGAACACGCAGCATGTTACATCACATTACGTCACCGTAAAAAAAAA
AATGTGAGAAAGACAAAAACTACCACTGGCCGTCTCAATTTCAAGATTATACACCACACTATTGCTTT
TTTTTTAATATAAATTAGCAGAAATCTCAAACGAATATTGGCAGGTTGGACTAAAAAAACTATCACTT
CAACAAGCATACACTTTGTTTTG

Aminoacid sequence of the *I. ricinus* PTHR2

MPSVFELVHKSAPLHIILADQDSNMQLSALQEAMLDCLKNDSSAVFDSGGLPTCPGVWDGILCWQPAP
AGTMAQVNCPGYVYGFDKQEMATKWCTENGTWYVSPEHGQPWTNYTRCFPQGATMPSENISIFEPHLP
TIKLISKVCYTVSLVTLLAAFLMLASMKRLRCPRNSLHMHLFISFILRALAFLLKDALFVDGIGLSTN
VDFDKENVDCKVFTSFWHYVLMANYCWILMEGLYLHSLVFLAFFTDSSGIIRYVALGWGLPVLFIIPW
VVVRATLEDTLCWTTNVRQDLFWIIRGPITASIVVNFVLFLNITRVLFAKLFASQTPQARKYRYRKWF
KSTLVLVPLFGAHYAFPLGMSPAAAGDLVELIWLYVDQLFSSSQGFVVALLYCFLNGEVQTELRKLFQ
RWRCDRRGPGCGDKQLQHRHSLFTQSVTFLSRGRSSIQSLHSVDRREGVAKSPSPNSIKSKMSQRLLS
NADQESLGAVKRMSIPEYQDPAKRTPEDKTGSQRKLNGGCPIRWECHDADSDGGAPADTSEDNLLEGV
IQKETCL*

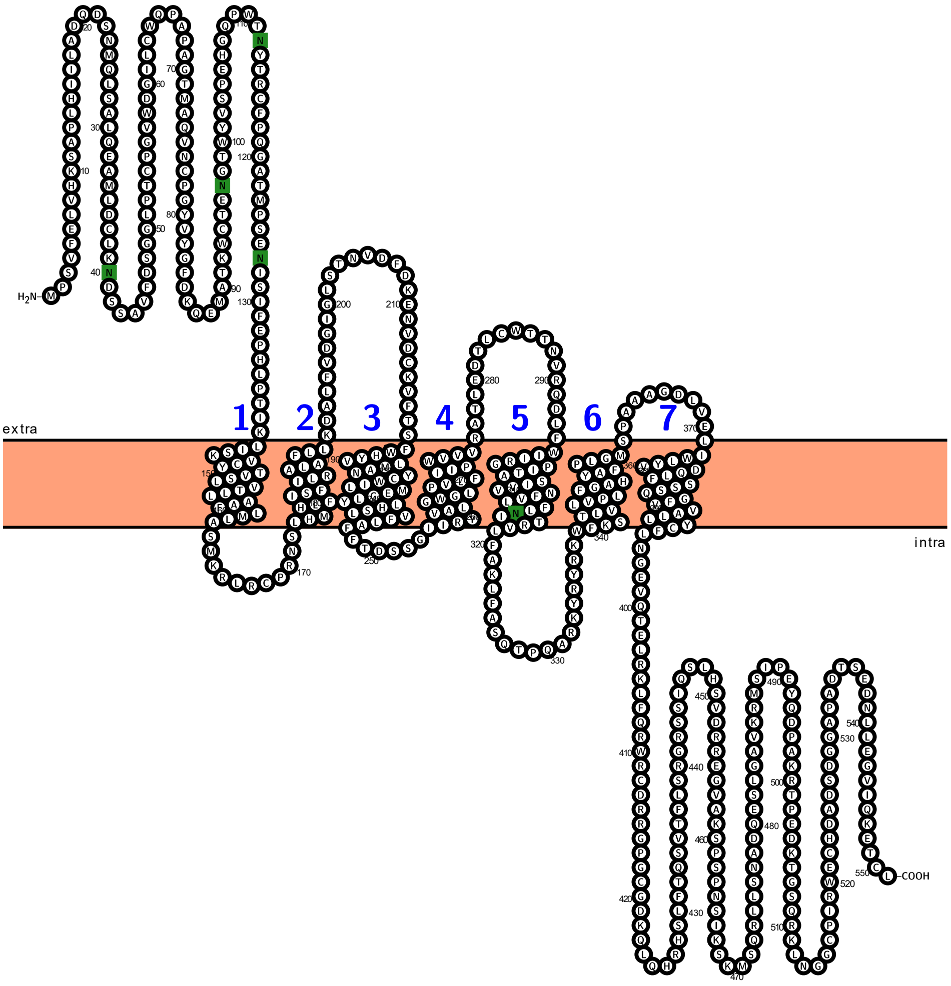


**Fig.2** Graphical visualization of *I. ricinus* PTHR2 made by Protter 1.0. Predicted N-glycosylation motif is marked green. Each of 7 transmembrane helices is numbered in blue.

```
Trica_PTHR1      ----------------------MSEFQDILNTAKRKCYASK-------------------    19
Trica_PTHR2      MTNI--------YVKKILDNQE-----TARKAAKEKCISMF----GPGVLQ----EFAEP    39
Iric_PTHR1       -----------------------MQTLSGLERAKRDCRVNESFTVDIGLSDESRLNLHD-    36
Iric_PTHR2       MPSVFELVHKSAPLHIILADQDSNMQLSALQEAMLDCLKND----------------SS-    43
                                               : *  .*

Trica_PTHR1      -TTEIITGKFCELIFDDILCWPPTPAGLLANQSCSNEYIKWTRNNYATRQCDSDGQWFIP    78
Trica_PTHR2      VYNETNKTLKCPAFWDTILCWPDTPSGTVINQSCPVYVAGFLASSNATRQCMENGSWYIR    99
Iric_PTHR1       IELNSTGGDFCPRVWDGILCWESAPADTTARAPCPEYLDGFDTTRTASKLCTENGTWYLS    96
Iric_PTHR2       AVFDSGGLPTCPGVWDGILCWQPAPAGTMAQVNCPGYVYGFDKQEMATKWCTENGTWYVS    103
                    :      *  .:* ****  :*:.   .  *      :     *:: * .:* *::

Trica_PTHR1      ENSTESWTNYSQCGNISEFYPILDDS-LRNHTLYNKWLPIIKNITQCGYILSTVSLIISL    137
Trica_PTHR2      D--NRTWSNYSSCYKDKITTIYLDLNKTNISGVLQNYAPVVQVISETGYIVSFATLIIAF    157
Iric_PTHR1       PLHGQTWTNYSQCFLAK---TS------DSLSMFEPHLPTIKLISKIGYTVSFVSLIVAF    147
Iric_PTHR2       PEHGQPWTNYTRCFPQGATMPS------ENISIFEPHLPTIKLISKVCYTVSLVTLLAAF    157
                     . *:**: *                   : :   * :: *::  * :* .:*: ::

Trica_PTHR1      FVFIRIKRLHCARNKLHIHLFASFVMRALMSLIKDGLFIEGTALPHEIIQINGKLVYNK-    196
Trica_PTHR2      AIMLFIKKLHCARNILHMHLFASFILRALTFIVIKSTFVEGLGLPSDLNYRNGSLYFDIN    217
Iric_PTHR1       IILASVKKLRCPRNSLHMHPFLSFILRALAVLVKDAIFVDGIGFSPNMDFS---------    198
Iric_PTHR2       LMLASMKRLRCPRNSLHMHLFISFILRALAFLLKDALFVDGIGLSTNVDFD---------    208
                  ::  :*:*:* ** **:* * **::***  :: .. *::* .:  ::

Trica_PTHR1      -TNFSWVCKAIISLWNYFIISNYMFLLMEGAYLHNLLFLKLL--SENGVVIYYSLGWGIP    253
Trica_PTHR2      SETNNWACKLLTSLWQYFITANYSWILMEGLYLHNLIFRALFADSSNSIKWYVVMGWGLP    277
Iric_PTHR1       --QDNVDCKVFTSFWHYVLMANYCWILMEGLYLHSLVFLAFFTLDSSGIIRYVALGWGLP    256
Iric_PTHR2       --KENVDCKVFTSFWHYVLMANYCWILMEGLYLHSLVFLAFFT-DSSGIIRYVALGWGLP    265
                     .  ** : *:*:*.: :** ::**** ***.*:*  ::  ....:  *  :***:*

Trica_PTHR1      LLFIIPWIVLKAGNENIYCWTTKSSKFIAMLIDVPIGLTVVINFILFTIIVRILFVKLTS    313
Trica_PTHR2      LIIVGFWVAARLLVEDNLCWTTHENYDVFLIIGIPTMVSILINLLLFMRISMVLYSKLRS    337
Iric_PTHR1       ILFLVPWVAVRAMLEDTYCWTTNDTKSYFWIIRGPITASIVVNFVLFINITRVLFVKLFT    316
Iric_PTHR2       VLFIIPWVVVRATLEDTLCWTTNVRQDLFWIIRGPITASIVVNFVLFLNITRVLFAKLFA    325
                 ::::  *:. :   *:  ****:       :*  *   ::::*::**  *  :*: ** :

Trica_PTHR1      MYI-QQRWTKYQKLIRAILILVPLFGIPYTISFVLSFYALEDQTFEIMWLFFDQTFTAFQ    372
Trica_PTHR2      PINED--SRRYQKWVKSTLVLVPLFGVHYALFLALYYLIKTNKIVEVVWLFCDLLFGSFQ    395
Iric_PTHR1       TQTHQARKYRYRKWFKSTLVLVPLFGAHYAFLLGMS-LAAAGDLLELVWLYIDQLFSSFQ    375
Iric_PTHR2       SQTPQARKYRYRKWFKSTLVLVPLFGAHYAFPLGMS-PAAAGDLVELIWLYVDQLFSSSQ    384
                     :    :*:* .:: *:******  *:: : :      .. .*::**: *  * : *

Trica_PTHR1      GLFASLVYCLLNSEVQMEIMRKYNSFKDRNREF-----------------KRRSRTISH-    414
Trica_PTHR2      GFFVAILYCFLNGEVKSEIQPHLYYFLTYLATNKYSKCLFPCRKKFLRSVVGRSSVCTTM    455
Iric_PTHR1       GFLVATLYCFLNAEVGAEARAAMARHRRTCCSDG-----LP-----LRVNKAQGSGCNEP    425
Iric_PTHR2       GFVVALLYCFLNGEVQTELRKLFQRWRCD----------------------RRGPGCGDK    422
                 *:..: :**:**.**  *                                  :.

Trica_PTHR1      ---------TQQIP----LTEELQEMPH---NA------GIKDQLCINKTSDYF*-----    446
Trica_PTHR2      S---CSSLYTNGVL-HRNSKCRLDSLPKIKTTD------KPKDHFCNNTSRRNSRSQRHS    505
Iric_PTHR1       --RARLSLLTRSLTLITRLEER-------------------------------------P    446
Iric_PTHR2       QLQHRHSLFTQSVTFLSRGRSSIQSLHSVDRREGVAKSPSPNSIKSKMSQRLLSNADQES    482
                          *. :

Trica_PTHR1      ------------------------------------------------------------    446
Trica_PTHR2      QAFNSSATIPTAET---------TLCIENNSGKCRSESEIN--------------EESLK    542
Iric_PTHR1       LSAKEDFDIPLHQVT*--------------------------------------------    461
Iric_PTHR2       LGAVKRMSIPEYQDPAKRTPEDKTGSQRKLNGGCPIRWECHDADSDGGAPADTSEDNLLE    542


Trica_PTHR1      ----------         446
Trica_PTHR2      MVVHSDF---         549
Iric_PTHR1       ----------         461
Iric_PTHR2       GVIQKETCL*         551
```

Multiple alignment of *I. ricinus* PTHRs with PTHR1 (XP_008192153.1) and PTHR2 (XP_015834182.1) of *T. castaneum*. Hormone receptor domain pfam02793 is marked magenta, and smart00008 is marked turquoise. Predicted transmembrane helices are marked gray. The conserved domains are derived from Conserved Domain Database tool of NCBI (Wang, et al. 2023).
